# Intradiscal Inflammatory Stimulation Induces Spinal Pain Behavior and Intervertebral Disc Degeneration *In Vivo*

**DOI:** 10.1101/2022.04.11.487751

**Authors:** Lauren E. Lisiewski, Hayley E. Jacobsen, Dan C. M. Viola, Hagar M. Kenawy, Daniel N. Kiridly, Nadeen O. Chahine

**Affiliations:** Department of Biomedical Engineering, Columbia University, New York, NY, United States; Department of Orthopedic Surgery, Columbia University, New York, NY, United States; Hughston Clinic Orthopedics, Nashville, Tennessee, United States

**Keywords:** intervertebral disc, inflammation, pain behavior, rat model, TLR4, spine pain

## Abstract

Degeneration of the intervertebral disc (IVD) is known to occur naturally over time, with the severity of pain varying widely. Other components of the degenerative environment, including structural disruption and inflammatory cytokine levels, and their correlation with pain severity have been studied. However, the role of the inflammatory environment in activating degenerative changes that manifest as a pain phenotype has not been elucidated. Previous studies have aimed to recreate the sustained inflammatory environment exhibited during human disc degeneration in a rat model. Most commonly, a puncture injury has been used causing structural damage and only initiating an acute inflammatory response. This study utilized injection of lipopolysaccharide (LPS), a pro-inflammatory stimulus, into the rat disc *in vivo* to create the desired sustained inflammatory environment independent of physical disruption. LPS injections resulted in upregulation of pro-inflammatory cytokines and an immunogenic response. The structural integrity of the IVD was also altered demonstrated by changes in histological score, disc height, and mechanical properties. Ultimately, a sustained inflammatory environment led to both local and radiating mechanical sensitivity, demonstrating that the pain phenotype experienced during disc degeneration can be initiated solely by a sustained inflammatory profile. Markers indicative of nerve ingrowth into the IVD were also expressed suggesting a potential mechanism for the pain exhibited by animals. This rat injury model will allow for future study of the direct relationship between inflammation and pain in the degenerative environment.

## Introduction

Low back pain is the leading cause of disability and is often associated with degeneration of the intervertebral disc (IVD)^1^. With aging, degeneration of the IVD occurs over time manifesting as extracellular matrix (ECM) disruption and ranging in pain severity from asymptomatic cases to severe pain^2^. Structural changes to the disc tissue are associated with pro-inflammatory and immunogenic responses as well as pathological innervation^3^. The inflammatory profile in human disc degeneration is characterized by sustained, elevated levels of TNFα and IL-1β. Alterations of the cytokine profile have been correlated with greater degrees of disc degeneration, with expression levels varying significantly between degenerated and herniated IVDs^4–6^. Interestingly, expression of cytokines TNFα and IL-1β have been shown to correlate with nerve growth factor (NGF) expression, a marker known to be expressed specifically in painful IVDs^7,8^. Additionally, degeneration grade and expression levels for markers indicative of innervation have also been correlated^9^. This suggests a causative relationship between inflammatory cytokines and the pain phenotype.

There is extensive evidence for the role of pro-inflammatory cytokines in certain types of disc disease. For example, herniated discs exhibit inflammatory cell infiltration, increased pro-inflammatory cytokines, as well as radiating pain symptoms due to mechanical compression and chemical irritation caused by the herniated tissue^10–13^. However, the role of inflammation in triggering localized pain of discogenic origin due to a contained but degenerated disc is less understood. Needle puncture or disc lesion injuries are the most popular type of models used to simulate disc degeneration *in vivo*, and have previously been implemented in both lumbar and caudal regions of mice, rats, and rabbits^14^. These studies have demonstrated that puncture or lesion successfully reproduces morphological changes associated with disc degeneration and behavior attributed to the experience of pain in animals. Both mechanical and thermal sensitivity were exhibited peaking at 3-9 months and 9-12 months, respectively^13,15–19^. However, inflammatory changes in these models are thought to be acute and transient post-injury, which differs from the presence of sustained chronic inflammation seen in human disc degeneration^20,21^. To assess the potential of disc inflammatory cytokines at triggering pain behavior, Lai et al. performed an intradiscal injection of TNFα, or other growth factors (VEGF and NGF) during needle puncture injury, and found accelerated expression of hind paw pain behavior compared to disc puncture alone^22^. Similarly, injection of TNFα in rat and porcine models resulted in decreased degeneration grade, with rat intradiscal injection leading to decreased paw withdrawal thresholds as well^23,24^. However, it remains unknown whether discal activation of inflammatory signaling can trigger localized spinal pain of discogenic origin.

Recent studies have highlighted the role of innate immune activation, particularly that of toll-like receptors (TLRs), in the pathogenesis of disc degeneration^25–27^. TLRs are members of a receptor family activated by damage associated molecular patterns (DAMPs) resulting in persistent pro-inflammatory signaling^28^. DAMPS such as high mobility group box 1 protein (HMGB1) have been shown to have degenerative effects in disc cells by signaling through TLRs and causing increased expression of inflammatory cytokines and matrix degrading enzymes^29^. Activation of TLR4 by LPS has been shown to induce a pro-inflammatory cascade in IVD cells ^30,31^. Further, injection of LPS directly into the NP space of rat caudal IVDs has caused moderate degenerative changes in the IVD, with increases in tissue levels of IL-1β, TNF-α, HMGB1, and macrophage migration inhibitory factor (MIF)^27^. However, the extent to which inflammatory signaling triggered by LPS injection contributes to pain behavior is unknown.

The goal of this study was to evaluate behavioral pain responses of rats after inflammatory activation of caudal IVDs by LPS injection *in vivo*. A range of LPS dosages were initially investigated to identify a dose causative of matrix integrity disruption. The behavior responses of rats to mechanical and thermal stimuli were evaluated locally in the spine and distally in the hind paw. Caudal IVDs, as opposed to lumbar, were selected to create an easily reproducible model with a simplified surgical procedure resulting in less complications. The objective of this study was to validate a new rat model of disc degeneration caused solely by inflammatory stimulation of the disc. Demonstration of structural changes, a sustained inflammatory profile, and the resultant local mechanical pain phenotype typical in human disc degeneration fills a critical gap in the area of animal models of disc degeneration. Additionally, this model will allow for future study of the mechanism of pain in the degenerating disc in relation to the inflammatory environment, independent of physical tissue disruption.

## Methods

Surgical procedures were performed separately without analgesia on 4 cohorts of animals, as described below. Institutional animal care and use committee approval was attained for each cohort prior to start of the experiments.

### Surgical Procedure – Part 1: Dose Response

Male Sprague Dawley rats (N=17) were anesthetized with isoflurane, and an incision was made exposing 4 caudal (C) motion segments, approximately C3-4 to C6-7. Phosphate buffered saline (PBS) or LPS was injected as previously described by Rajan et al^27^. Briefly, A 33G needle was used as it is <10% of average rat caudal disc height minimizing physical tissue disruption^32^. It was inserted 4mm into the center of the disc with clamp guidance. 2.5 μl of saline, as a sham control, or LPS at a single dose (either 1, 10, or 100 μg/mL) were injected slowly over 60 seconds into alternating discs using a microliter syringe. The incision was closed with 4-0 nylon sutures and animals were allowed unrestricted activity before being euthanized at 2, 7, or 28 days post-injury.

### Surgical Procedure – Part 2 & 3

Male Sprague Dawley rats (Part 2: N=9, Part 3: N=10) were anesthetized with isoflurane, and an incision was made in the caudal spine to expose 3 caudal motion segments, approximately C4-5 to C6-7. A33G needle was inserted 4mm into the center of the disc with clamp guidance. 2.5 μl of saline, as a sham control, or LPS (100 μg/mL) were injected slowly over 60 seconds into the caudal discs of each animal using a microliter syringe, with all discs in an animal receiving the same treatment. Animals were euthanized at 28 days after injury.

### Surgical Procedure – Part 4

Male Sprague Dawley rats (N=12) were anesthetized with isoflurane. Using fluoroscopic guidance, caudal motion segments C5-6 to C8-9 were identified and exposed. Sutures were used to mark above and below the IVDs of interest. 2.5 μl of saline or LPS (100 μg/mL) was injected into all discs of each animal as described above. Animals were euthanized at 14 or 28 days post-injury.

### Dimensions and Biomechanical Testing

Bone-disc-bone motion segments were collected from part 1 and the surrounding connective tissue was removed. Motion segment height and diameter were measured with digital calipers. Using the measured values, cross sectional area (CSA) and aspect ratio were calculated using the following formulas:

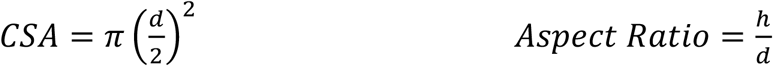

where d represents diameter and h represents sample height. Data was normalized to the corresponding sham from the same animal, and unpaired t-tests between LPS and sham were used to determine statistical significance.

For mechanical testing, motion segments were subjected to unconfined compression between 2 stainless steel platens while submerged in PBS. Using the Instron testing frame (Instron 5566) equipped with 10 and 100 N load cells, samples were first pre-loaded with a 0.1 N tare load followed by cyclic loading to 3 N applied at 0.1 Hz for 30 cycles. Disc segments were then subjected to a creep load of 3 N until equilibration. The resulting displacement was measured and creep strain was calculated. The creep modulus and dynamic modulus were then calculated using the following formulas:

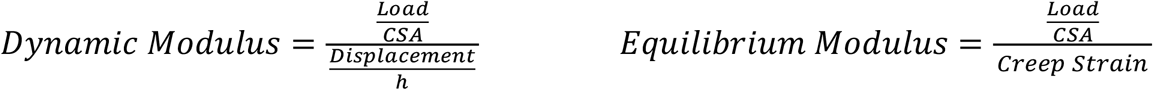

Statistical significance between LPS and sham groups was determined using unpaired t-tests. Biomechanical differences between the 14 and 28 day time points were also investigated.

### Biochemical Content

Individual discs were dissected after biomechanical testing, and separated into annulus fibrosis (AF) and nucleus pulposus (NP) using a biopsy punch. Tissue wet weights were measured, samples were dried in vacuum desiccator, and then dry weights were measured. Water content was calculated based on percent difference in wet and dry weights. Samples were then digested overnight in papain (Sigma-Aldrich). Tissue digests were analyzed for DNA content using a Picogreen assay (Invitrogen), GAG content using the Blyscan assay (BioVendor), and collagen content using the orthohydroxyproline (OHP) assay^33^. Biochemical content was reported as sample concentration of biochemical compound normalized to wet weight. Statistical significance was calculated using one-way ANOVAs with a Fisher’s LSD post-hoc test for the comparison between LPS doses and sham groups.

### Behavioral Testing

Behavioral testing was performed on rats from part 2 and 3 described above. Animals in part 2 studies underwent the Pressure Algometry Measurement (PAM) test in the dorsal tail, Von Frey test in the hind paw, and tail flick test in the distal tail. Animals in part 3 studies underwent the PAM test in the dorsal tail, Von Frey test in the ventral trail, Hargreaves tests in the tail base, and tail flick test in the distal tail. Behaviors were evaluated at day 0, as a baseline pre-injury measurement, and then at 1, 3, 7, 14, and 28 days post-injury. Testing was conducted in a quiet room and animals were acclimated to each test’s equipment and the experimenter. The group assignment of the animals was not known to the experimenter at the time of testing.

Mechanical hyperalgesia was measured using the Von Frey and PAM tests. The Von Frey test was performed either in the hind paw or the ventral tail^16^. Rats were placed in individual cubicles on top of a suspended wire mesh surface. The animals were allowed to acclimate to the testing space for 15-20 minutes. Von Frey filaments (Ugo Basile, Semmes-Weinstein set of monofilaments, cat. 37450) were presented perpendicularly to the plantar surface of the selected hind paw (part 2 studies) or ventrally to the base of the tail (part 3 studies), where they were held in position for approximately 2-3 seconds with enough force to cause a slight bend in the filament. Testing was started with a 2 gram (g) force filament and decreased or increased to a maximum force of 15 g based upon the response from the rat or lack thereof. Positive responses included an obvious withdrawal of the hind paw or tail from the filament and/or flinching behaviors and licking. Data was collected until three positive responses or the maximum force threshold was reached without achieving an observable response. The average of the forces resulting in a positive response of the maximum force value was recorded and normalized to the baseline average in each group. The PAM procedure was adapted from Kim et al. and was conducted using an Ugo Basile instrument (cat. 38500) accompanied by a large transducer (cat. 58500-2)^34^. The rat was manually restrained with a glove and/or wrapped in a towel, and allowed to acclimate to the holder until resistance ceased. The PAM transducer was placed on the testers thumb and slowly pressed on the base of the tail directly on the dorsal skin over the experimental discs. The force was slowly increased at 100 g/second until an audible vocalization or obvious physical response was observed. Following response, the associated force was recorded. A cut off of 1,500 g was set to avoid tissue damage. Recorded forces were normalized to the baseline average within group for each study. After normalization, part 2 and part 3 study data were plotted together.

Thermal hyperalgesia was measured using the tail flick and Hargreaves tests according to the methods described by Mohd Isa et al^16^. The tail flick test was conducted using the Ugo Basile instrument (cat. 37360). Rats were previously acclimated to the handler with no immediate acclimation necessary. The rat was manually restrained by the holder and/or wrapped in a towel to hold the rat steady on the top of the instrument. Radiant heat was focused 4-7 cm from the distal end of the tail. Heat was applied until the rat responded by flicking the tail away from the heat source and the response time was recorded. The infrared (IR) intensity was set at 20 and a cut off time was set at 15 seconds to prevent tissue damage. For the Hargreaves test, rats were places in the Hargreaves arena (Ugo Basile) and allowed to acclimate for 15-20 minutes. Heat was applied ventrally at the base of the tail. Response time was recorded as the withdrawal time. A positive response was considered to be withdrawing, flinching, licking, or biting. IR intensity was set to 40 with a cutoff time of 20 seconds. In all behavioral tests a two-way ANOVA was used with group (Sham vs. LPS) and time as variables. Between group differences at each time point and change in behavior relative to baseline within each group were evaluated using a Fisher’s LSD post-hoc test.

### Disc Height Measurement and Histological Analysis

Fluoroscopic disc height measurements and histological analysis were performed on disc segments from rats in part 4 studies. Fluoroscopic images were taken of all IVDs of interest pre-injury, and 14 or 28 days post-injury. Three measurements of each disc height (D1-D3) and the adjacent vertebral body heights (V1-3, V4-6) were used to calculate the disc height index (DHI) for each disc level using the following formula:

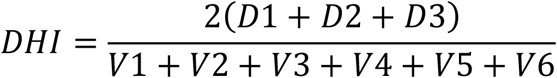

Post-injury DHI values were normalized to each disc’s pre-injury DHI to elucidate changes in disc height between groups and over time. Statistical significance was determined using a one-way ANOVA with a Tukey post-hoc test for all comparisons.

For histological analysis, 14 or 28 days post-procedure disc segments were dissected and submerged in 4% paraformaldehyde (PFA) for 48 hours. Following fixation with PFA, disc segments were washed 3 times with PBS and decalcified in 14% ethylenediaminetetraacetic acid (EDTA) at 4°C for approximately 2 weeks. Discs were again washed with PBS and transferred to 70% ethanol for transportation to the Molecular Pathology Shared Resource (MPSR) facility at the Columbia University Herbert Irving Comprehensive Cancer Center (HICCC) where paraffin embedding, sectioning, and Safranin O-Fast Green staining was performed. Sections were imaged and histological grading was conducted using a consensus grading scale for rat IVDs detailed by Lai et al^35^. Eight categories were scored from 0 to 2 by two graders with 0 being healthy and 2 being degenerated, and average scores for each category were determined. Overall histological grade out of 16 was calculated for each disc with statistical significance determined using a two-way ANOVA with a Šídák post-hoc test between conditions and time points. The distribution of histological grades in each subcategory was plotted and significance of average scores between sham and LPS groups was calculated using an unpaired t-test.

### Gene Expression Analysis

Gene expression analysis was completed on IVDs from part 4 rats 14 or 28 days after injury. Discs were isolated from the caudal spine with the AF and NP manually separated using a scalpel. Each region was individually snap frozen in cryogenic vials. Tissue was pulverized using a bead homogenizer and cells were lysed with TRIzol and chloroform prior to phase separation. A high salt solution in combination with isopropanol was utilized for precipitation of RNA before purification with the RNeasy Mini Kit (Qiagen) according to the manufacturer’s protocol. Expression of genes was measured using RT-qPCR to evaluate changes in macrophage markers (iNOS, Arg1), inflammatory cytokines (TNFα, IL-1β), and nociceptive markers (CGRP, NGF, BDNF). Primer sequences can be found in Table 1.

**Table 1.**
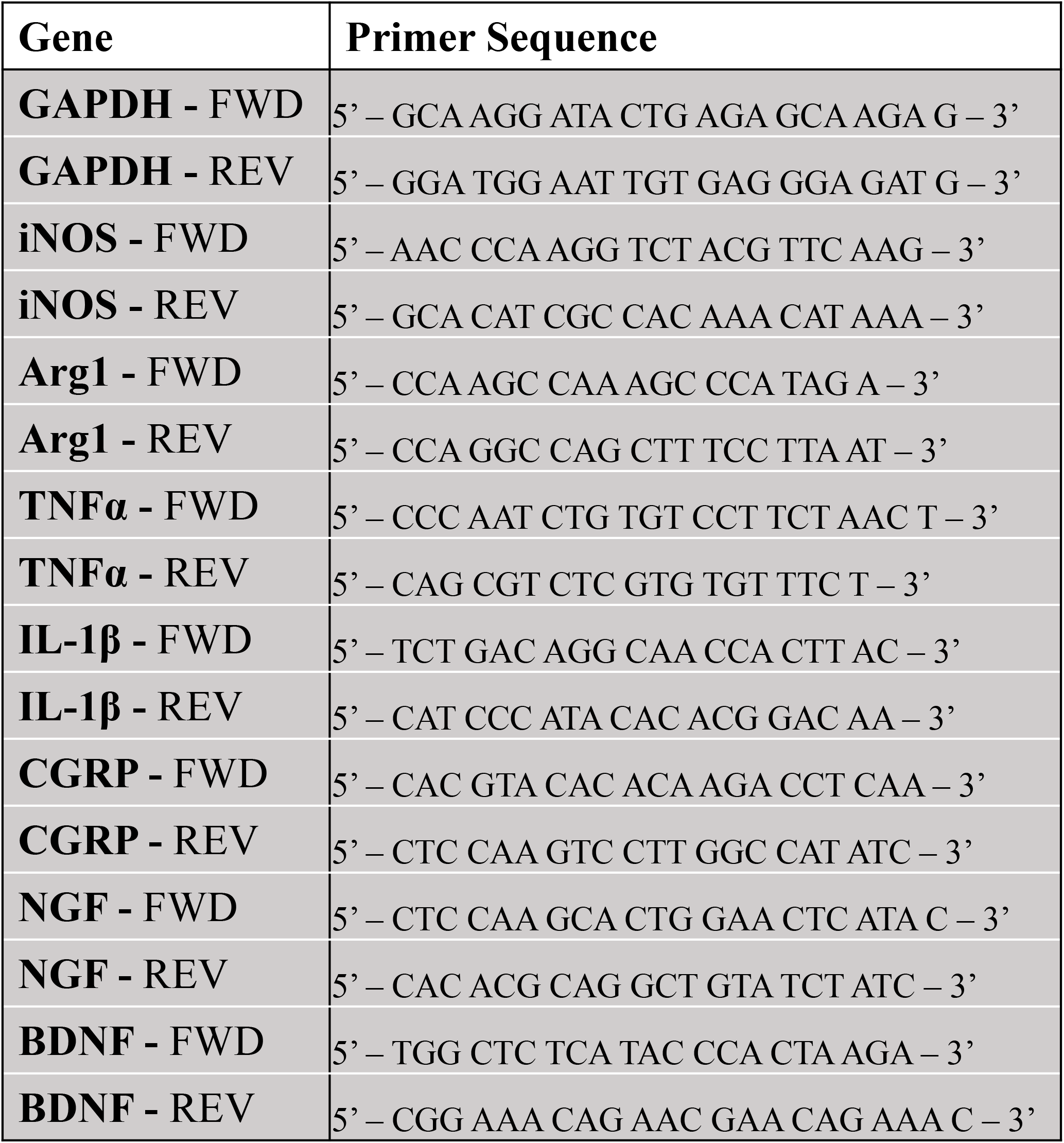
Primer Sequences

## Results

### LPS Dose Response: Dimensions, Biomechanics, and Biochemical Composition (Part 1)

On Day 2 post injection, no significant differences were observed in the cross-sectional area and aspect ratio of samples injected with any of the LPS doses compared to those injected with saline. On day 7, a dose of 1 or 10 µg/mL did not result in dimensional or biomechanical changes of motion segments compared to sham. However, there was a significant increase (p<0.05) in cross sectional area and a significant decrease (p<0.05) in aspect ratio in disc segments injected with 100 µg/mL LPS vs. sham. The sham group was shown as a dashed line for graphical representation of the change in dimensional measurements (Figure 1A-B). There was also a trend for decreasing equilibrium modulus (p=0.07) and dynamic modulus (p=0.06) in 100 µg/mL LPS group compared to sham at day 7. Similar decreases in both mechanical properties (equilibrium modulus: p=0.08; dynamic modulus p=0.07) were observed at day 28 post-injection. Additionally, time dependent decreases in equilibrium modulus were observed at day 28 compared to day 7 in the sham (p<0.05) and 10 µg/mL LPS (p<0.05) groups (Figure 1C-D).

**Figure 1.**
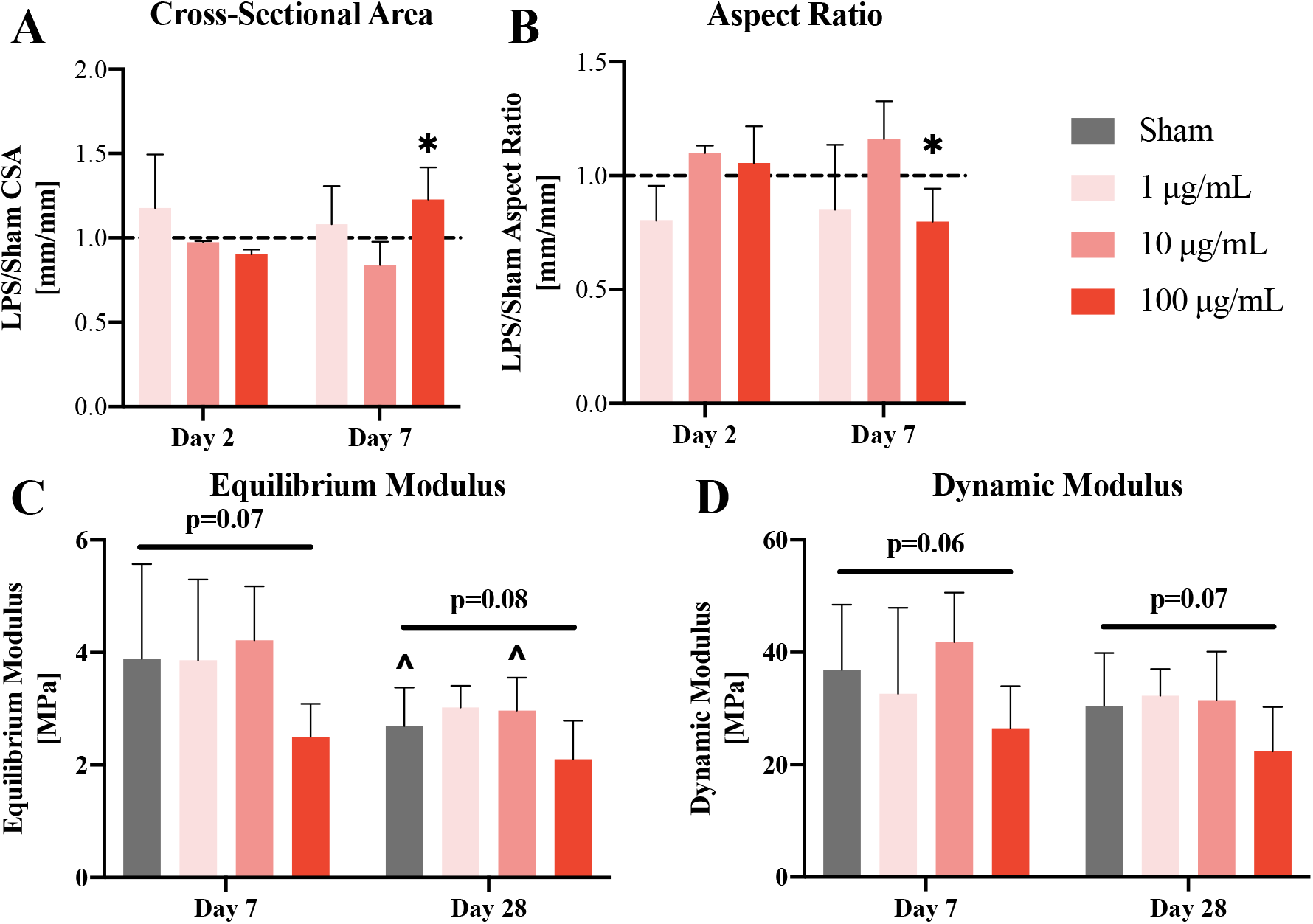
Dose Response (Dimensions and Mechanical Testing) **(A)** Cross sectional area and **(B)** Aspect ratio of intervertebral discs after injection of LPS normalized to sham. * p<0.05 LPS injection in comparison to normalized sham value indicated by dashed line. **(C)** Equilibrium modulus and **(D)** Dynamic modulus measurements for sham discs injected with saline or LPS. ^ p<0.05 day 28 in comparison to day 7, p-value for comparison between sham and 100 µg/mL.

In the AF, DNA content significantly increased (p<0.05) at day 7 post-injection with 10 µg/mL LPS in comparison to sham, while other LPS doses had comparable levels to sham. No significant differences were observed in the DNA content of the NP in all dose groups (Figure 2A). AF collagen content was similar to sham 7 days post-LPS injection. However, in the NP, a significant increase in collagen content was observed in the 100 µg/mL LPS group compared to sham (p<0.01) (Figure 2B). At day 7, GAG content in the AF significantly increased (p<0.05) in the 100 µg/mL LPS group compared to sham, but not in the 1 and 10 µg/mL LPS groups (Figure 2C). No significant differences in NP or AF GAG content were observed at day 28 (Figure 2E). Water content of AF and NP of all groups at day 7 and day 28 showed no significant differences (Figure 2D,F).

**Figure 2.**
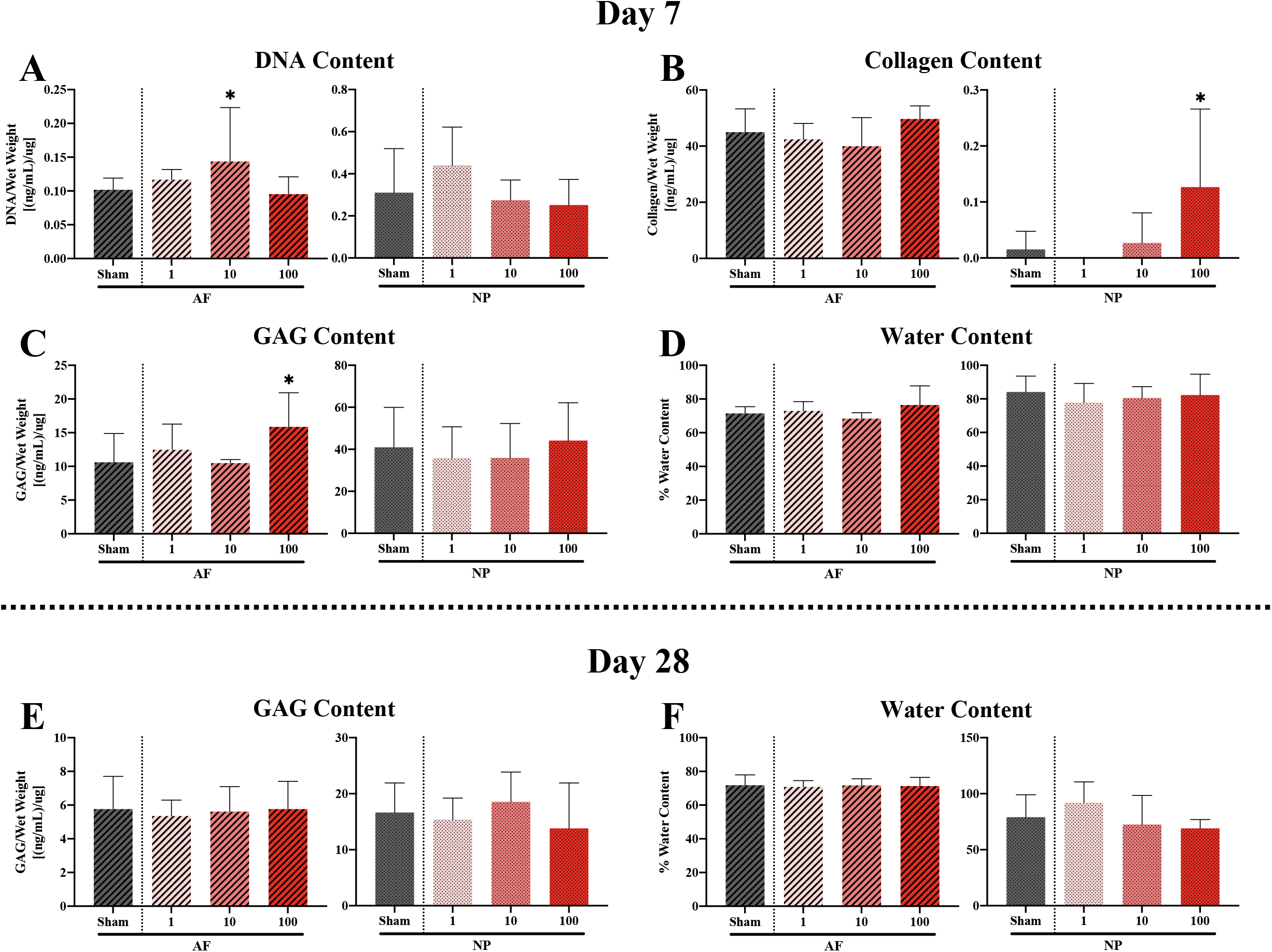
Dose Response (Biochemistry) **(A)** DNA content, **(B)** Collagen content, **(C)** GAG content, and **(D)** Water content of IVDs 7 days after injection with saline or LPS. **(E)** GAG content and **(F)** Water content of discs 28 days after injection. * p<0.05 LPS injection in comparison to sham.

### Behavior Testing: Mechanical and Thermal Sensitivity (Part 2 & 3)

The PAM test in dorsal tail and Von Frey tests in ventral tail and hind paw were used to measure changes in mechanical sensitivity longitudinally over 28 days in comparison to baseline, and between the LPS and sham groups at each time point. After LPS injection, the PAM test resulted in a significantly lower withdrawal time at day 14 (p<0.05) and 28 (p<0.05) with a trend observed at day 21 (p=0.07) in comparison to sham. Withdrawal time for both sham and LPS decreased within 1 day of surgery, and then increased over time; however, magnitudes remained significantly different from baseline up to day 28 in both groups (Figure 3A). In the Von Frey test of the ventral tail, withdrawal force decreased significantly compared to baseline at day 1 through day 28 in the LPS group, but not in the sham group (Figure 3B). In the hind paw, withdrawal force was similar to baseline up to day 7 in both groups. In the LPS group, the withdrawal force at day 7 was significantly decreased (p<0.05) compared to baseline, while in the sham group significance (p<0.0001) from baseline was observed at day 14 (Figure 3C). At day 21, the withdrawal force in the LPS group was significantly lower than in sham (p<0.05).

**Figure 3.**
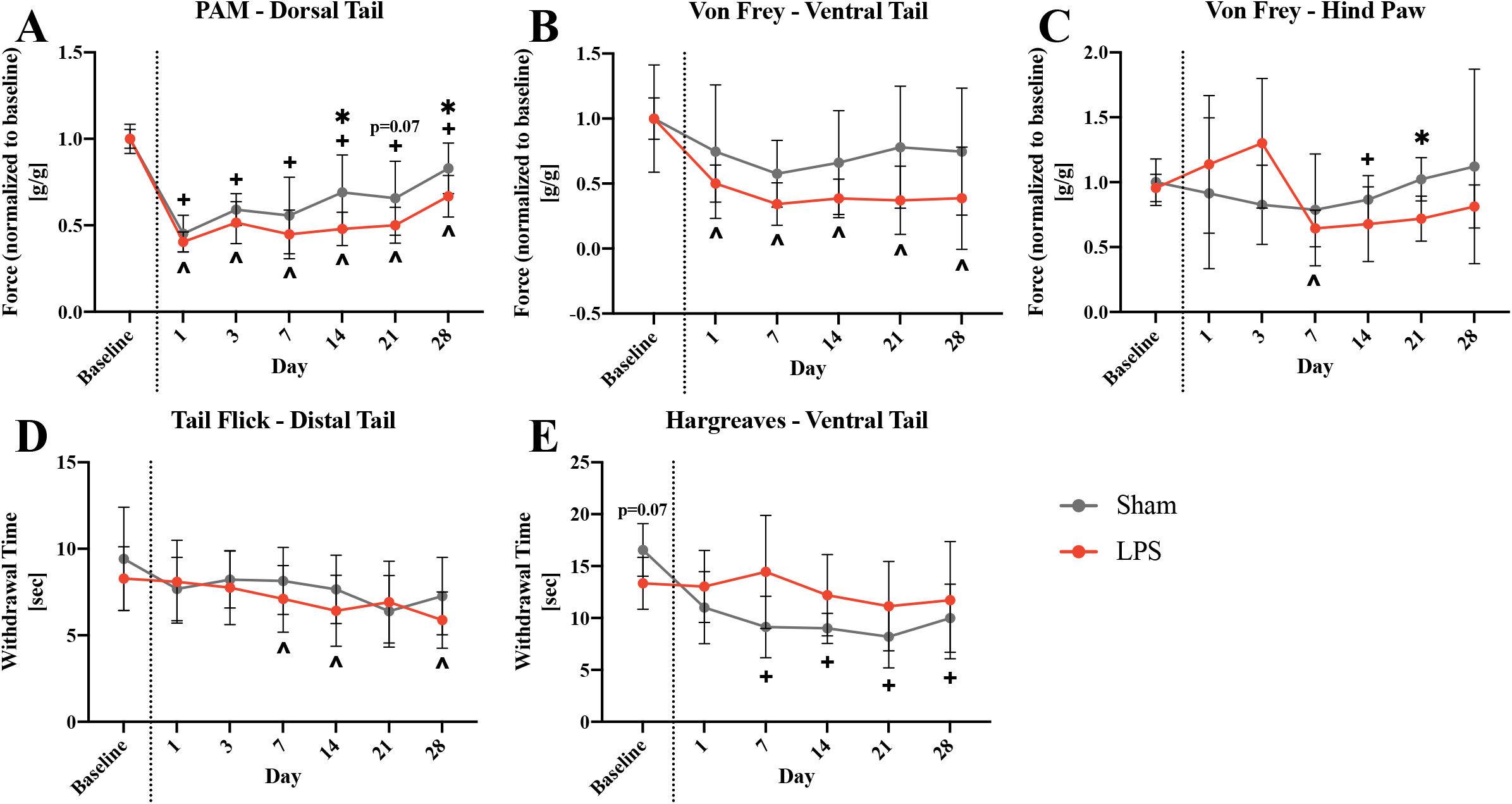
Behavior Tests. **(A-C)** Behavioral tests measuring mechanical sensitivity at baseline and longitudinally 1 to 28 days after injection of saline or LPS. **(D-E)** Behavioral tests measuring thermal sensitivity at baseline and longitudinally 1 to 28 days after injection. * p<0.05 for LPS in comparison to sham (p-value shown for trends defined as 0.05<p<0.1), ^ p<0.05 LPS significantly different from baseline at indicated timepoint, + p<0.05 Sham significantly different from baseline at indicated timepoint.

The tail flick and Hargreaves behavioral tests were utilized for measurement of changes in thermal sensitivity after LPS injection. Neither thermal sensitivity test resulted in significant differences between the sham and LPS injection groups. In the tail flick test of the distal tail, significantly lower withdrawal time was measured at days 7 (p<0.05), 14 (p<0.01), and 28 (p<0.05) after LPS injection in comparison to baseline. No differences compared to baseline were observed in the sham group. The Hargreaves test of the ventral tail resulted in a significant decrease in withdrawal time from day 7 to day 28 (all p<0.01) compared to baseline in the sham group. Differences in comparison to baseline were not observed in the LPS group. However, a trend (p=0.07) for higher withdrawal time in sham group compared to LPS at baseline was observed.

### Histology: Structural and Morphological Changes (Part 4)

At days 14 and 28 post-injection, fluoroscopic images of all injected IVDs were analyzed in comparison to pre-injection images. Disc height index normalized to pre-injection exhibited a significant decrease (p<0.001) in the LPS group over time from day 14 to day 28, and in comparison to sham at day 28 (p<0.01) (Figure 4B). Representative images of the most degenerated and least degenerated discs from each group and time point are presented to illustrate the heterogeneity in the histological changes (Figure 4A). Total histological score was significantly higher (p<0.05) in the LPS group compared to sham groups 28 days post-injection (Figure 4C). Of the 8 structural variables evaluated, 5 categories exhibited significantly different average scores between sham and LPS at day 28 including NP shape (p<0.01), NP area (p<0.05), NP-AF border appearance (p<0.01), AF lamellar organization (p<0.01), and AF tears/fissures/disruptions (p<0.05). A trend towards lower histological scores in the endplate of the LPS group was observed in comparison to sham (p=0.08). No significant differences in NP cell number and NP cell clustering and morphology were observed at day 28. Additionally, no significant differences were observed at day 14 in any category of the histological grading scale (Figures 4D-K).

**Figure 4.**
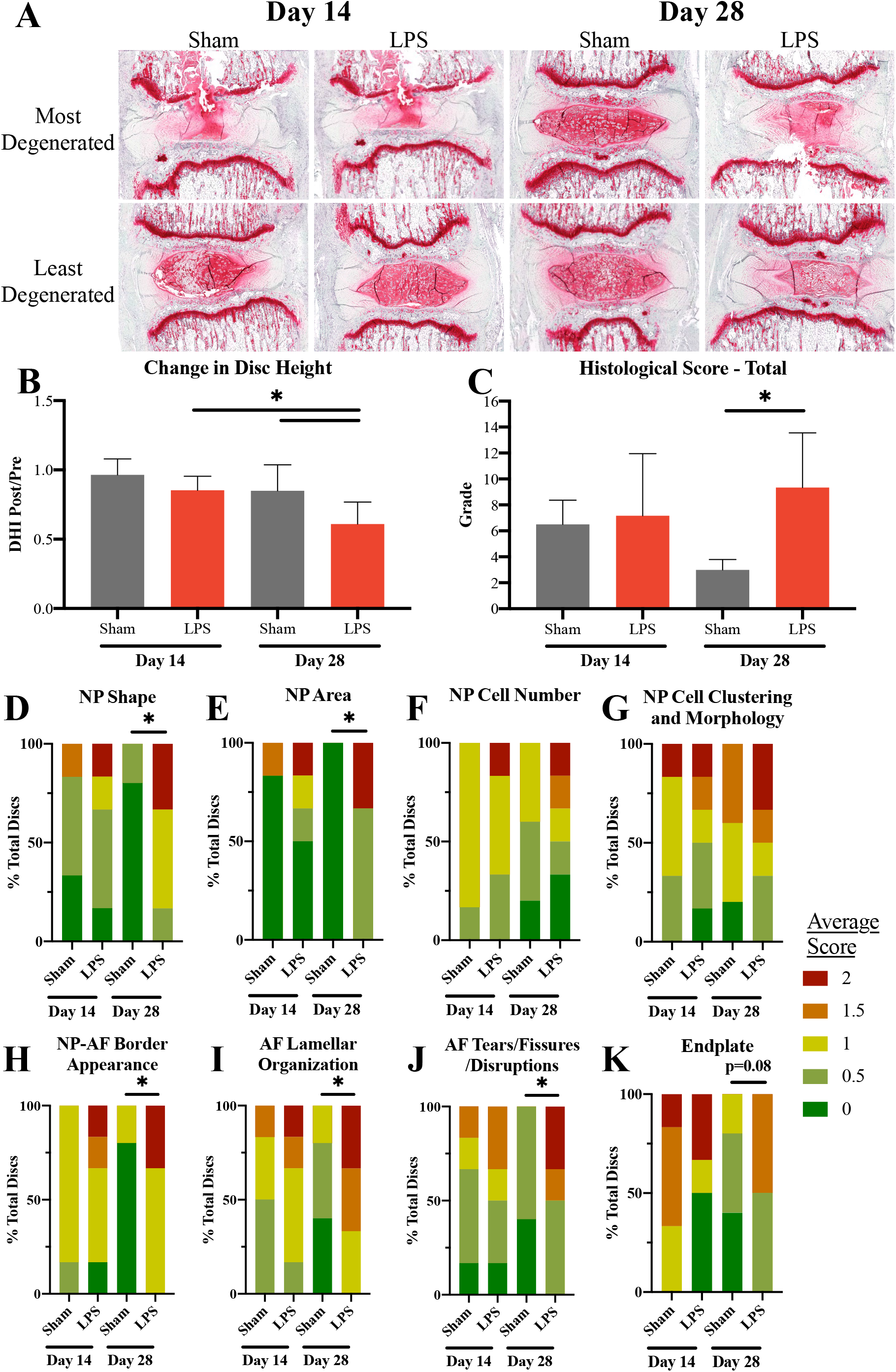
Histology. **(A)** Representative images of most and least degenerated IVDs from sham and LPS injected groups at 14 and 28 days after injection with saline or LPS. **(B)** Change in fluoroscopically measured disc height post-injection normalized to pre-injection height. **(C)** Total histological score of sham and LPS injected discs. **(D-K)** Histological score categorical breakdown and score distributions. * p<0.05 for LPS in comparison to sham (p-value shown for trends defined as 0.05<p<0.1)

### Gene Expression: Macrophage, Inflammatory, and Nociceptive Markers (Part 4)

AF and NP tissue regions were analyzed separately at day 14 and day 28 for gene expression of markers related to macrophages, inflammatory cytokines, and nociception. Expression of the pro-inflammatory macrophage marker, iNOS, was significantly upregulated in AF of LPS injected groups at day 14 (p<0.05) and day 28 (p<0.05). In the NP, iNOS expression also significantly increased in the LPS group compared to sham at day 14 (p<0.01). Expression of the anti-inflammatory macrophage marker, Arg1, trended lower in the NP of the LPS group compared to sham at day 14 (p=0.08); however, this trend reversed 28 days post-injection showing a trend towards increased (p=0.09) expression in the NP (Figure 5B). No significant changes in Arg1 were found in the AF (Figure 5A). Expression of the pro-inflammatory cytokines, TNFα and IL-1β, were significantly upregulated in AF of the LPS group in comparison to sham at day 28 (TNFα: p<0.05; IL-1β: p<0.01), with a significant increase (p<0.05) in TNFα over time from day 14 to day 28 (Figure 5C). In the NP, a significant increase in IL-1β was observed at day 14 in LPS vs. sham (p<0.01), but no difference in TNFα expression was observed. Differences in expression for either inflammatory cytokine were not seen in the NP at day 28 (Figure 5D).The expression of the nociceptive markers CGRP, NGF, and BDNF increased in the LPS group in comparison to sham in both AF and NP at day 14 and day 28. In the AF, BDNF expression in the LPS group was trending higher at day 14 (p=0.07), while CGRP and NGF expression were similar to sham. In the NP at day 14, CGRP and BDNF expression significantly increased in the LPS group compared to sham (CGRP: p<0.05; BDNF: p<0.001), while NGF exhibited a trend for higher expression in LPS vs. sham (p=0.06). At day 28, all three nociceptive markers were significantly upregulated in the AF of the LPS group in comparison to sham (CGRP: p<0.01; NGF: p<0.01; BDNF: p<0.05), and expression of CGRP at day 28 was significantly greater (p<0.01) than at day 14 (Figure 5E). In the NP, only CGRP remained significantly increased (p<0.05) in comparison to sham at day 28 (Figure 5F).

**Figure 5.**
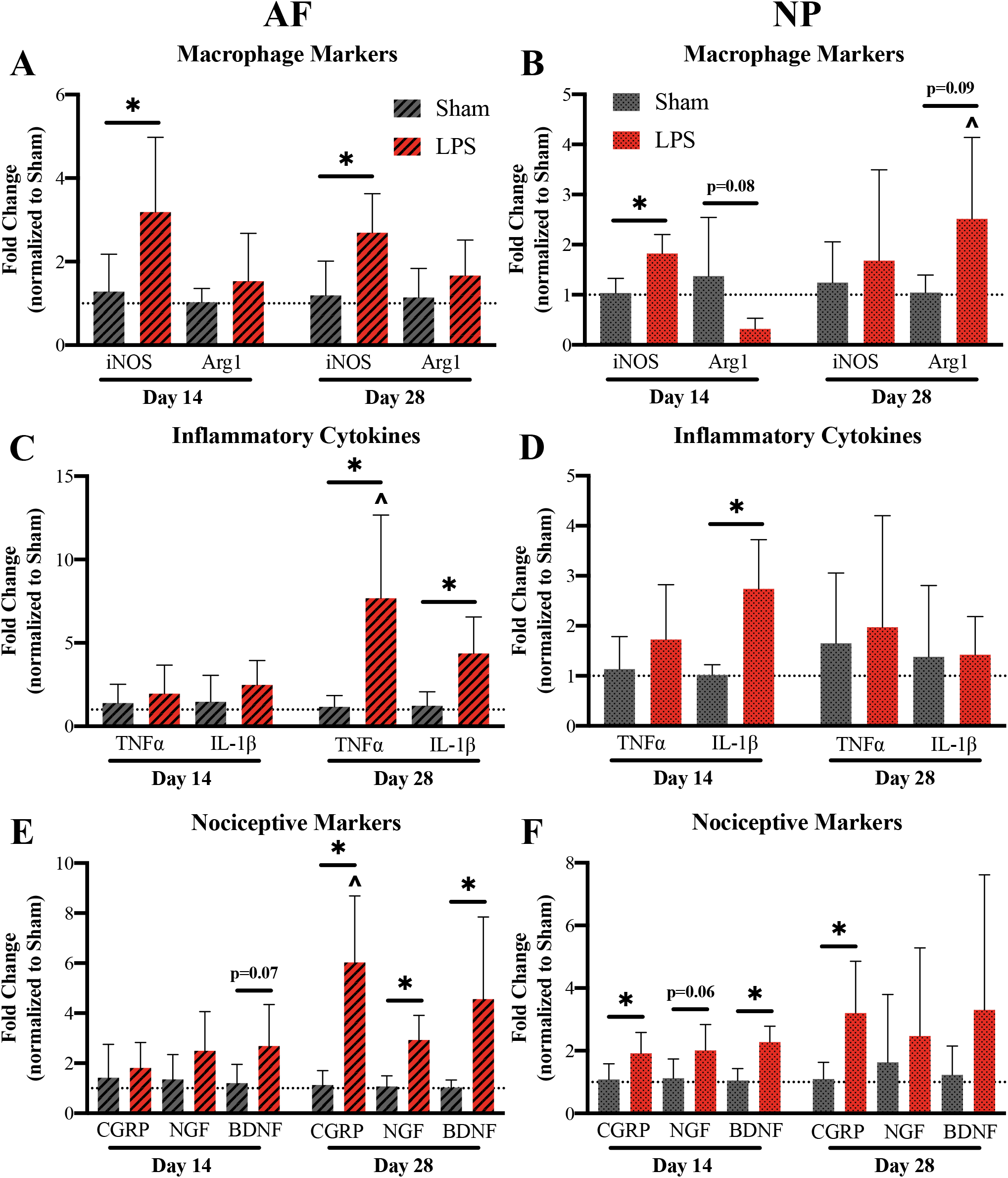
Gene Expression. Gene expression for macrophage markers in **(A)** AF and **(B)** NP tissue. Measurement of expression of inflammatory genes in **(C)** AF and **(D)** NP tissue. Expression of markers for nociceptive genes in **(E)** AF and **(F)** NP tissue. * p<0.05 or p= listed value LPS in comparison to sham, ^ p<0.05 Day 28 in comparison to day 14.

## Discussion

This study focused on the development of a disc degeneration model in rat caused solely by inflammatory stimulation of the IVD using LPS injection into the NP space. Injection of LPS triggered an inflammatory and immunogenic response indicated by increased expression of markers for macrophages and inflammatory cytokines. The inflammatory cascade caused structural and biomechanical changes in the disc, as well as biochemical alterations indicative of fibrosis. Rats that underwent LPS injection demonstrated a pain phenotype, correlating with the increase in nociceptive markers expressed by disc tissue.

### LPS Activation of TLRs Promotes Inflammatory Signaling

Recent studies have focused on the activation of TLRs during disc degeneration and their role in propagating the production of inflammatory cytokines. LPS is well-known for its role in the activation of TLR4, which is known to be expressed on the surface of IVD cells and correlate with degeneration grade^25^. When stimulated with LPS *in vitro*, NP cells have shown upregulation of gene and protein levels of TLR4 followed by the inflammatory cytokines, TNFα and IL-1β^26,27,30^. This study supports these findings demonstrating increases in gene expression of TNFα and IL-1β after LPS injection (Figure 5C-D).

Importantly, the increased expression for markers of inflammatory cytokines is sustained for up to 28 days, although the tissue location of the inflammatory upregulation varies over time.Similar findings of variable timelines for expression of each gene were demonstrated by Rajan et al. showing an overall increase in inflammatory cytokine expression after *in vivo* LPS injection at day 1, followed by a decrease in TNFα at day 7 while IL-1β continued to increase^27^. Additionally, Ponnappan et al. showed an increase in IL-1β expression in the NP, but not in the AF 10 days after cytokine stimulation in an explant model, agreeing with our results^36^. The maintenance of the inflammatory environment over a long period of time is critical to validation of this injury model, as it is reflective of what is experienced during human disc degeneration.

Other injury models, mostly utilizing the puncture injury method, have attempted to recreate this sustained inflammatory environment. Ulrich et al. utilized multiple puncture injuries over a period of 6 days, while Lai et al. incorporated an additional injection of TNFα during puncture to produce the desired results^21,22^. The injury model used in this study is the first to demonstrate a sustained inflammatory profile resulting in a pain phenotype without additional physical disruption to the tissue.

### Degenerative Environment Leads to Matrix Breakdown Independent of Tissue Disruption

It is often thought that the structural integrity of the disc is only disrupted from frequent mechanical overloading or injury^37^. Previous studies have validated this method for promotion of degeneration demonstrating decreased disc height and increased histological score after puncture injury^13,15,17,19^. However, we demonstrated similar degenerative changes in the structural integrity of the disc stimulated only by exposure to an inflammatory stimulus. At 28 days post-LPS injection, a significant decrease in disc height as well as total histological score was recorded (Figure 4A-C). Interesting, when histological score is analyzed by subcategory, it suggests that the majority of degenerative changes involved the ECM of both the AF and NP, while the endplate as well as NP cell number and cell morphology remain unchanged (Figure 4D-K).

Extracellular matrix disruption is a common measure of degeneration severity, with a general trend towards decreasing collagen with age^2^. It has also been demonstrated that human NP and AF cells treated with IL-1β express decreased levels of collagen and aggrecan^4,38^. These results are indicative of the imperative role that cytokines play in the process of ECM degradation during IVD degeneration. This process has been further investigated *in vivo*, showing increased aggrecanase expression after injection of LPS or IL-1 without physical disruption of the tissue structure^27,31^. Our findings do not agree with previous studies that demonstrated decreased collagen and aggrecan content after mechanical injury or inflammatory stimulation in a rat model. Although, biomechanical testing results did show a significant loss of compressive mechanical properties of the IVD in dynamic loading and at equilibrium after LPS injection, indicating the softer material that would result from the biochemical changes seen in previous literature (Figure 1C-D). Contrarily, our results demonstrated minimal changes in GAG and water content and an increase in collagen content of the NP after LPS injection, instead agreeing with the common understanding that the IVD becomes fibrotic during aging and degeneration (Figure 2B)^39^.

### Cytokines Encourage Immune Cell Recruitment

The IVD is considered an immune-privileged structure due to its lack of both blood vessels and nerves, with these features existing only in the outer AF in the healthy state. When this structural integrity is compromised, blood vessel grow into the disc from adjacent vertebral bodies or the damaged AF allowing macrophages to enter^3,40^. Recruitment in this way is further motivated by production of cytokines and chemokines by NP cells^38,41^. Macrophages can be broadly categorized as either M1, pro-inflammatory macrophages, or M2, anti-inflammatory macrophages. Each subtype plays a different role in the degenerative environment and is therefore recruited at differing times in the degenerative cascade. This behavior was demonstrated by Nakawaki et al. through increased expression of M1 macrophage markers in the IVD through day 14, followed by a delayed increase in M2 macrophage markers from day 7 to day 28 after injury^42^. These findings are in agreement with our study, that demonstrated a greater initial increase of the M1 marker, iNOS, at day 14 followed by an increase in the M2 marker, Arg1, at day 28 (Figure 5A-B). Additionally, other studies have shown that the increase in inflammatory cytokines, TNFα and IL-1β, seen after injury, are decreased when macrophages are depleted^43^. This supports macrophages contribution to the production of cytokines and the critical role that they play in perpetuating the inflammatory environment during disc degeneration^41^.

### Pain Behavior Stimulated by the Degenerative Environment

Pain is a common occurrence during disc degeneration and is often related to the ingrowth of nerves into the disc. It is believed that nerves enter the disc through structural disruption where there are less proteoglycans present^40^. However, even when nerves are present, they do not always result in pain. Symptomatic disc degeneration is comprised of two important components: nerve ingrowth into the IVD and sensitization of those nerves^44^. Previous studies have looked at the progression of innervation in a mouse IVD after injury, and how this manifests as pain behavior. They demonstrated that mechanical sensitivity did not peak until 3-9 months after injury^13^. This behavior could explain why our model showed trends towards differences in the mechanical hyperalgesia tests, PAM and Von Frey, but resulted in inconsistent statistical significance (Figure 3A-C). Additionally, an initial thermal sensitivity was predicted to be due to exposure of nerves to NP contents released during injury^45^. However, after this initial response, it did not reappear until 9-12 months after puncture injury, with most innervation located in the dorsal aspect prior to 6 months after injury^13,40^. Our results are in agreement with these previous studies, as the rats showed limited responsiveness to thermal stimuli up to 28 days after LPS injection, especially in the ventral tail (Figure 3D-E). Perhaps a longer timeframe may have better encompassed the entirety of inflammatory effects on pain.

Nonetheless, inflammatory cytokines are known to play a critical role in the induction of pain, with greater cytokine levels being correlated with higher degrees of pain^46,47^. Indeed, an injection of the cytokine TNFα was shown to increase degeneration grade and promote a pain phenotype in comparison to an injection of PBS. However, when anti-TNFα was injected instead, both of these effects were mitigated^23^. These findings implicate cytokines in the sensitization of nerve fibers. It has also been demonstrated that treatment of IVD cells with TNFα or IL-1β stimulates production of NGF^42^. This supports the involvement of cytokines in nerve ingrowth independent of the pain phenotype as well. Our findings demonstrate a similar upregulation in the expression of NGF after inflammatory stimulation using LPS injection. Additionally, CGRP is known to be involved in sensitization of nerves with increased expression being shown after injury^48^. The results of this study confirm this finding, demonstrating increased expression of CGRP at day 28 in both the NP and AF (Figure 5E-F). Nerve ingrowth stimulated by NGF and sensitized by CGRP provides a possible mechanism for the pain phenotype demonstrated in this rat model after LPS injection.

### Limitations

The limitations of this study include the short timeframe of 28 days, the use of only male rats, and the lack of analysis of how gene expression translates to protein expression in this model. The chosen timeframe of 28 days is an extension of those used in some previous studies; however, in long-term models over a period of 12 months, degeneration grade continued to change over time and some pain behaviors reemerged at later time points^13,40^. The use of only male rats is not ideal because gender specific pain behaviors and IVD biomechanics have been previously demonstrated^17^. Finally, gene expression results do not always directly translate to protein production so it will be important to validate that expression in future studies.

## Conclusion

This study demonstrated an immunogenic response, structural changes, and pain behavior as a result of direct inflammatory stimulation of the IVD *in vivo*. Taken together, these findings indicate that inflammation alone, in the absence of traumatic IVD disruption, triggers the degenerative cascade. Future studies will utilize this rat injury model to delineate the role that inflammation plays in the development of discogenic pain and the related mechanisms at play during disc degeneration.

## Acknowledgements

The authors acknowledge Jansher Khan for assistance with studies and thank Dr. Huan Yang for assistance with training on the behavior tests. Histology services were provided by the Molecular Pathology Shared Resource (MPSR) facility at the Columbia University Herbert Irving Comprehensive Cancer Center (HICCC). This research was funded in part by NIH R01 AR069668 and R01 AR077760.

