## Supplemental Figures 1 & 2 for "Intradiscal Inflammatory Stimulation Induces Spinal Pain Behavior and Intervertebral Disc Degeneration *In Vivo*"

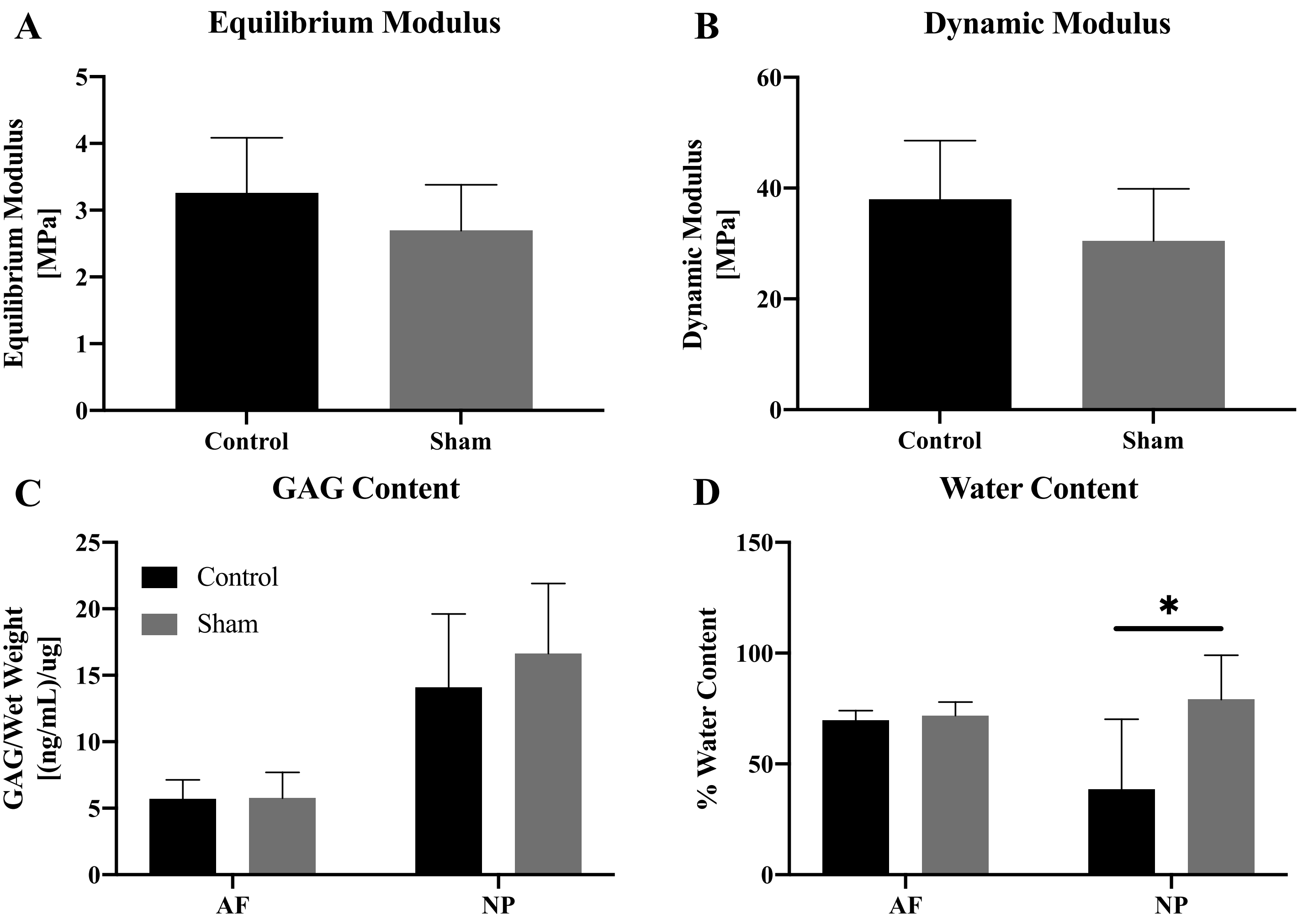


**Supplementary Figure 1. (A)** Equilibrium modulus and **(B)** Dynamic modulus measurements for rats with incision only (control) or incision with saline injection (sham). **(C)** GAG content and **(D)** Water content of IVDs 28 days after incision only or incision with saline injection. * $p<0.05$ control in comparison to sham. Statistical significance determined using unpaired t-test.

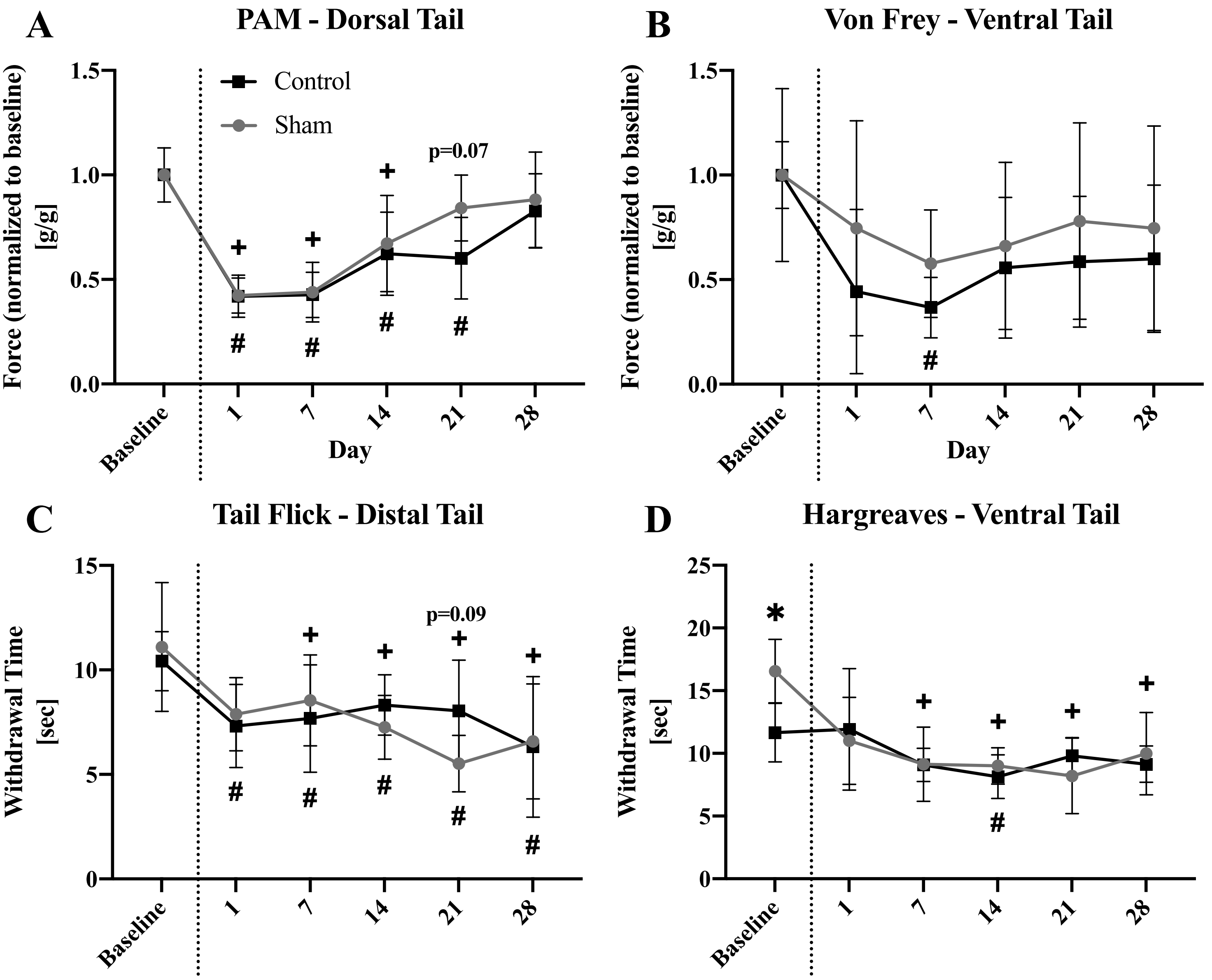


**Supplementary Figure 2. (A-B)** Behavioral tests measuring mechanical sensitivity at baseline and longitudinally 1 to 28 days after incision only (control) or incision with saline injection (sham). **(C-D)** Behavioral tests measuring thermal sensitivity at baseline and longitudinally 1 to 28 days after injection. * $p<0.05$ for incision only in comparison to sham (p-value shown for trends defined as $0.05<p<0.1$), # $p<0.05$ Control significantly different from baseline at indicated timepoint, + $p<0.05$ Sham significantly different from baseline at indicated timepoint. Statistical significance determined using 2-way ANOVA.
